# Functional consequences of pre- and postsynaptic expression of synaptic plasticity

**DOI:** 10.1101/075317

**Authors:** Rui Ponte Costa, Beatriz E.P. Mizusaki, P. Jesper Sjöström, Mark C. W. van Rossum

## Abstract

Growing experimental evidence shows that both homeostatic and Hebbian synaptic plasticity can be expressed presynaptically as well as postsynaptically. In this review, we start by discussing this evidence and methods used to determine expression loci. Next, we discuss functional consequences of this diversity in pre- and postsynaptic expression of both homeostatic and Hebbian synaptic plasticity. In particular, we explore the functional consequences of a biologically tuned model of pre- and postsynaptically expressed spike-timing-dependent plasticity complemented with postsynaptic homeostatic control. The pre- and postsynaptic expression in this model predicts 1) more reliable receptive fields and sensory perception, 2) rapid recovery of forgotten information (memory savings) and 3) reduced response latencies, compared to a model with postsynaptic expression only. Finally we discuss open questions that will require a considerable research effort to better elucidate how the specific locus of expression of homeostatic and Hebbian plasticity alters synaptic and network computations.

## Introduction

Synapses shape the computations of the nervous system. The combination of thousands of excitatory and inhibitory synaptic inputs determine whether a neuron fires or not. Furthermore, the synapse is known to be a key site of information storage in the brain, although not the only one [1]. Changes in the synapses are hypothesized to allow neuronal networks to change function and to adapt through Hebbian and Hebbian-like mechanisms. At the same time, large perturbations in activity levels such as those occurring during synaptogenesis or eye-opening require negative feedback so that the network can keep its activity level within reasonable bounds and continue performing its computational tasks properly [2, 3]. Such homeostatic control of neuronal activity can occur through changes in intrinsic neuronal properties such as control of dendrite excitability [4, 5], somatic excitability [6, 1] and movement of the axon hillock relative to the soma [7]. However, in this review we focus on homeostatic processes at the synapse such as synaptic scaling, which provides a form of negative feedback to counter changes in the activity levels, while providing synaptic normalisation and competition among inputs [8, 9].

As we explain in detail in this review, irrespective of whether synaptic plasticity is Hebbian or homeostatic, the expression locus of plasticity matters. A fundamental distinction is whether the change is pre- or postsynaptic. Changes in the number of postsynaptic receptors typically only modify the synaptic gain. However, long-term changes in the presynaptic release probability alter the short-term dynamics of the synapse [10, 11, 12, 13, 14, 15, 16]. Synaptic dynamics such as short-term depression and facilitation describe how the synaptic efficacy changes during repeated stimulation of the synapse over a time course of hundreds of milliseconds [13, 17, 18, 19]. These short-term modifications of synaptic efficacy (reviewed in [19]) have been proposed to underlie computations like gain control [20], redundancy reduction [21] and adaptive filtering [22]. In the context of a recurrent neuronal network, they can affect the activity dynamics and allow the formation and switching among attractor states [23, 24], and have been proposed as the basis for working memory [25].

Synaptic plasticity can thus affect network dynamics, but this poses several questions: What are the functional implications of expressing long-term plasticity pre- or postsynaptically? What are the underlying expression mechanisms? Why is there such a large diversity in the expression? And why is there sometimes both pre- and postsynaptic expression? In this review, we begin by discussing pre- and postsynaptic components of Hebbian and homeostatic synaptic plasticity. Then we examine some of the consequences of the variability of the expression locus of synaptic plasticity, including those that we recently identified using a biologically tuned computational model of neocortical spike-timing-dependent plasticity (STDP) [16].

## The biological underpinnings of pre- and postsynaptic expression of plasticity

As old as the field of long-term synaptic plasticity itself is the question of how precisely information is stored in neuronal circuits. Historically, Donald Hebb and Jerzy Konorski argued for the strengthening of already existing connections between neurons as a means for information storage, whereas Santiago Ramon y Cajal favoured the growth of new connections [26]. Several relatively recent studies have found evidence that the formation of new synapses is important for long-term information storage in neuronal circuits [27, 28, 29, 30]. Indeed, there is strong evidence both in mammals and in the sea slug Aplysia that structural plasticity via formation of new afferent inputs is essential for protein-synthesis dependent long-term memories [31]. The creation of new afferents would correspond to an increase in the number of release sites (see Box 1: Methods), but it should be noted that the number of release sites might be different from the number of anatomical contacts [e.g. 32].

### BOX1

#### Methods to determine the locus of plasticity

The properties of synaptic release can be used to determine the locus of synaptic plasticity by a variety of methods. Among these there are methods for studying vesicle release, such as FM1-43 dye labelling to explore changes presynaptic release [67], glutamate uncaging to explore changes in postsynaptic responsiveness or spine size [68, 69], measuring NMDA:AMPA ratio to look for insertion of postsynaptic receptors [70, 48], employing the use-dependent NMDA receptor blocker MK-801 to look for changes in glutamate release [71, 72], or exploring changes in paired-pulse ratio suggesting a change in probability of release [15, 48] (although see [73]).

It is also common to employ spontaneous release as a metric of the locus of expression, as each spontaneously released vesicle gives rise to a well-defined single postsynaptic quantal response known as a miniPSC. This approach is often used in studies of homeostatic plasticity (e.g. [74]), because here it is important to measure synaptic changes globally across a majority of inputs to a cell, but this method has also been used to explore Hebbian plasticity [75, 70]. An increase in miniPSC frequency in the absence of a change in miniPSC amplitude is typically interpreted as indicating higher release probability or an increase in the number of synaptic contacts, while an increased miniPSC amplitude is most often thought to reflect an increase in postsynaptic responsiveness due to more efficacious postsynaptic receptors. Alternative interpretations of spontaneous release experiments are, however, also possible, for example in the case of AMPA-fication of silent synapses, which leads to an apparent change in release probability even though unsilencing is a postsynaptic process [75].

In the scenario where individual synapses are monitored, it is possible to employ methods that rely on the response variability. One such method is non-stationary noise analysis [76], which has been used to determine the effect of homeostasis on inhibitory connections [77], although this method can be unreliable for dendritic synapses [78]. In the related coefficient of variation (CV) analysis, the peak synaptic response is modelled as a binomial process. The process has as parameters the release probability Pr, and the response to each vesicle, the quantal amplitude *q*. These parameters are assumed identical across the N release sites, and indeed such coordination has been found [65]. The CV — which is experimentally quantified as the response standard deviation over the mean — is independent of q, namely 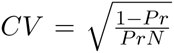, and therefore an increase in the mean without an increase in CV can be interpreted as a postsynaptic increase of *q* [79]. Conversely, if plasticity is presynaptically expressed, then a change in CV is expected, since the CV is a measure of noise and since the chief source of noise in neurotransmission is the presynaptic stochasticity of vesicle release. The CV analysis method does, however, come with several caveats. In particular, accidental loss or gain of afferent fibers in extracellular stimulation experiments, or unsilencing or growth of new synapses will confuse the results [79]. It is also not obvious that release is independent at different sites, in which case the binomial model is not suitable [79]. By assuming that one of the parameters does not change during the experiment (e.g. fixed N as is reasonable to assume in some plasticity experiments [80, 81]) the variance and mean of postsynaptic responses can be used to estimate 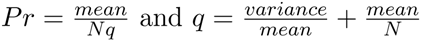 [33, 82, 16].

An alternative way to determine whether synaptic changes correspond to alterations of release probability or of quantal response amplitude is to examine the postsynaptic response to a pair or a train of presynaptic stimuli. The idea is that when the release probability is high, the vesicle pool will be depleted more quickly, leading to a more strongly depressing train of postsynaptic responses. When combined with CV analysis, this method can be used to measure all three parameters — *Pr, N*, and *q* — of the binomial release model [83]. By fitting these phenomenological models before and after plasticity induction, one can determine which combination of parameters were changed due to plasticity. It should be noted that experimental results from paired-pulse experiments should also be treated with caution. For example, unsilencing or specific postsynaptic upregulation of release sites with quite different release probability may lead to changes in short-term dynamics that could erroneously be interpreted as presynaptic in origin, even though the actual site of expression is postsynaptic [73]. There are also postsynaptic contributions to synaptic short-term dynamics [84, 85, 86], that can complicate the interpretation of experiments. It is therefore better to employ several methods in parallel in the same study — such as CV analysis, paired-pulse ratio, NMDA:AMPA ratio, and spontaneous release [15, 48] — to independently verify the locus of expression.

Recently, inference methods of short-term plasticity and quantal parameters have been introduced [87, 88, 89]. The sampling method of [87] is particularly well suited to deal with the strong correlation and uncertainty in the synapse parameters. Based on this method we revealed interesting variations between different neuronal connections and proposed more informative experimental protocols based on irregular spike-trains, which would be promising to apply in plasticity experiments.

With already existing connections between neurons, there are essentially only two possible ways of increasing synaptic strength: either presynaptic release is increased, or postsynaptic receptor channels are upregulated [33, 34]. Both can be achieved in a number of ways. The presynaptic release probability is controlled by various factors, such as the number and sensitivity of presynaptic calcium channels, as well as other presynaptic ion channels that can modulate neurotransmitter release (such as the epithelial sodium channel ENaC in case of synaptic scaling at the Drosophila neuromuscular junction [35, 36]), the setpoint of presynaptic calcium sensors involved in eliciting neurotransmitter release, e.g. the synaptotagmins 1, 2 and 9 [37], and the size of the pool of readily releasable vesicles as well as its replenishment rate (in case of homeostasis, see [38, 39]) [13, 37].

The postsynaptic contribution to the synaptic response is determined by the number and location of postsynaptic receptors, as well as their properties (e.g. conformational state [40] and subunit composition [41, 42]). In addition, the geometry of the extracellular space and the apposition of the release sites have also been suggested as important determinants of the response amplitude [43, 44].

Experimentally, determination of the expression locus is far from trivial and a battery of techniques has been applied (see Box 1). In long-term potentiation (LTP) experiments, evidence for most of the above mechanisms has been found. The historic pre versus post controversy is now typically interpreted as a reflection of the diversity of LTP phenomena, which we now know depends on multiple factors such as age, synapse state, neuromodulation, synapse type, and induction protocol [33, 45, 46, 47, 48, 49, 50, 51, 52] (but see [53]). A combination of pre- and postsynaptic expression is also possible [33].

A similar pre- or postsynaptic expression question exists for synaptic homeostasis. While most studies have focused on postsynaptic expression, also here a wide variety in expression, including presynaptic expression [54, 55, 56], has been observed, and for instance whether the expression is pre- or postsynaptic appears to depend on developmental stage [57, 58]. Sometimes diversity in mechanisms can even be observed within one system. For instance, in homeostatic plasticity experiments in the hippocampus both pre- an postsynaptic expression was observed, while some CA3-CA3 connections were unexpectedly *reduced* after activity deprivation, other connections strengthened as expected, perhaps to prevent network instability [59]. Also some forms of synaptic scaling at the Drosophila and mammalian neuromuscular junction (NMJ) are presynaptic: loss of postsynaptic receptors is compensated by increased transmitter release, which restores the mean amplitude of evoked EPSPs [36, 60]. A presynaptic locus of expression of homeostatic plasticity at the NMJ is perhaps to be expected, given that the postsynaptic partner — the muscle myotube — does not integrate its inputs like a neuron does, but rather serves to fire in response to activation at the synaptic input. The pre- and postsynaptic components of the NMJ are therefore tightly co-regulated in synaptogenesis and after damage to ensure proper activation of the muscle [61], so when post- synaptic NMJ sensitivity is reduced, it is in this context not entirely surprising that the presynaptic machinery compensates accordingly by upscaling neurotransmitter release. This example illustrates how the locus of expression must be understood in the context of function of the synapse type at hand.

Further indication that the exact expression locus is functionally important comes from the fact that the expression of both short-term plasticity [62] and long-term plasticity [52] can depend on pre- and post-synaptic cell-type. In the case of short-term plasticity, connections from the same presynaptic neurons onto different cells can short-term depress or facilitate depending on the target cell type [63, 64], while multiple connection between two neurons are often highly similar [65]. Similarly, while spike-timing-dependent plasticity (STDP) exists at both horizontal and vertical excitatory inputs to visual cortex layer-2/3 pyramidal cells, the mechanistic underpinnings as well as the precise temporal requirements for induction are different [66]. Such specificity suggests that the specific locus of expression of long-term plasticity at a given synapse type is meaningful for the proper functioning of microcircuits in the brain, as otherwise tight regulation of expression locus would not have arisen during the evolution of the brain.

## Pre- and postsynaptic expression of STDP

While the diverse pathways of plasticity induction and expression are increasingly unravelled, their functional roles are still largely an open question. Recently, we have started exploring some of these consequences using computational models of STDP. In STDP experiments, where spikes from the presynaptic neuron are paired with millisecond precision with postsynaptic ones, the question of pre- versus postsynaptic expression has been extensively examined as well. Depending on factors such as synapse type, brain area and experimental conditions, there is evidence for both pre- and postsynaptic changes [15, 48, 90, 91, 66, 92]. Because of the synapse-type specificity of STDP [52], we used STDP data of connections between visual cortex layer-5 pyramidal cells only [93, 15, 48]. At this synapse it has been observed that using STDP induction protocols potentiation has both pre- and postsynaptic components [48], while LTD is expressed presynaptically only [15]. Presynaptic-only time-dependent LTD has also been found in other synapse-types and brain areas [90, 92].

Our model of STDP allows for distinct pre- and postsynaptic expression, Fig.1a. This phenomenological model relies on three dynamic variables, one which tracks past presynaptic activity *x*_+_(*t*), and two that track postsynaptic activity, *y*_+_(*t*) and *y*_−_(*t*). These traces increase with every spike and decay exponentially between spikes. The plasticity is expressed as a function of the traces, but in contrast to traditional STDP models where just the synaptic weight changes as a function of them [94], here both the release probability and the quantal amplitude are independently modified. In our model, we assume that the number of release sites *N* is fixed and that it does not change on the time-scale of the experiments, consistent with experimental observations [80, 81]. However, the model could be straightforwardly generalised to also include changes in *N*.

**Figure 1:**
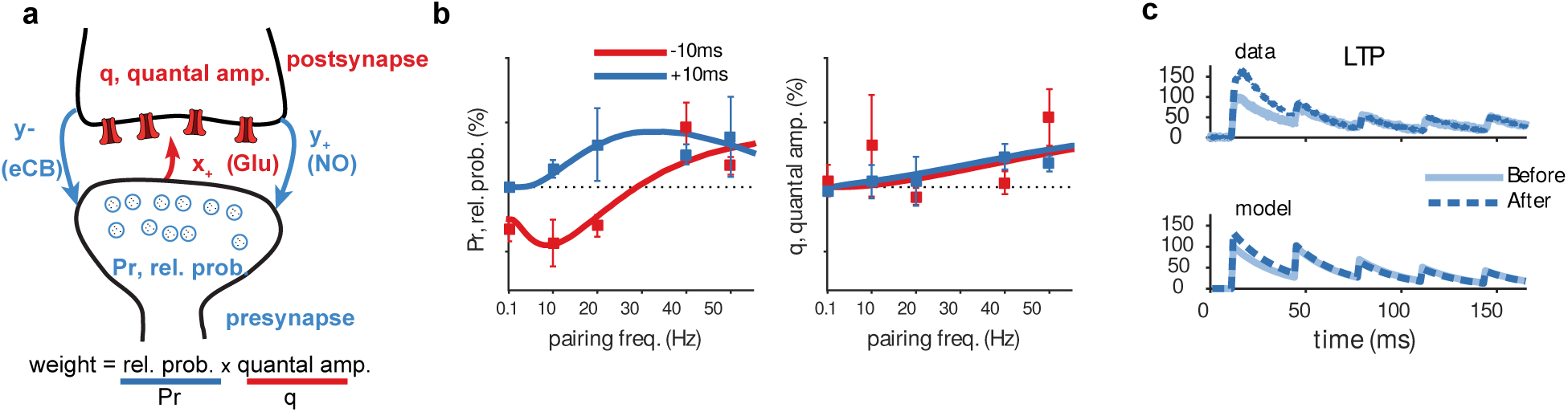
A schematic of our biologically tuned STDP model with pre- and postsynaptic expression. a) The synaptic weight is the product of the release probability P and the quantal amplitude *q*. Changes in these parameters due to STDP are modelled as functions of presynaptic activity trace *x*_+_ and postsynaptic activity traces *y*_+_ and *y*_−_. b) The fitted model captures the estimated changes in release probability (left) and quantal amplitude (right) for both positive timing (presynaptic spikes 10 ms before postsynaptic ones; blue) and negative timing (presynaptic spikes 10 ms after postsynaptic ones; red), as a function of the frequency of STDP pairings. Symbols indicate data, while lines denote the model fit. c) After LTP, the release probability is enhanced, which leads to stronger short-term depression. The change in short-term synaptic dynamics in the model (bottom) mimics the data (top). Panels b and c are reproduced from [16].

Even though we model the observed phenomenology rather than the biophysical or mechanistic details, with caution the components of the model can be interpreted to correspond specific physiological components. The presynaptic trace (*x*_+_), for example, could represent glutamate binding to postsynaptic NMDA receptors, which when depolarised by postsynaptic spikes unblocks NMDA receptors, leading to classical postsynaptic LTP [34]. Similarly, the postsynaptic trace *y*_+_ can be interpreted as retrograde nitric oxide (NO) signalling, which is read out by presynaptic spikes and leads to presynaptically expressed LTP [48]. Finally, the postsynaptic trace *y*_−_ can be linked to endocannabinoid (eCB) retrograde release, which triggers presynaptically expressed LTD when coincident with presynaptic spikes [15, 90, 92].

As mentioned above, we fitted our model to experimental data of one synapse type only (layer-5 pyramidal cells onto layer-5 pyramidal cells in the visual cortex) [93, 15, 48], across different frequencies and timings. To ensure the biological realism of the model, we further constrained the model fitting by using data from NO and eCB pharmacological blockade experiments in which either presynaptic LTD or LTP expression alone was abolished [48]. Furthermore, we verified that our model captured the expected interaction of short and long-term plasticity correctly (see Fig.1c), which permits the exploration of the functional implications of changes in short-dynamics due to the induction of long-term plasticity.

In the current model neither LTD nor LTP depend on the state of the synapse - the values of *q* and *Pr.* As a result the current model does not have a (non-trivial) fixed point, and as the fitting to the data only considered the *relative* changes in these parameters, the initial conditions were arbitrarily set to *q* = 1. An improved model could include state dependence in the plasticity to 1) create a fixed point and a realistic weight distribution, and 2) allow fitting to data that takes into account that plasticity might depend on the state (see also Discussion). Such extensions would however require more data. Similarly it might be possible to model plasticity at the level of voltage[95] or even calcium [96] to capture finer details observed experimentally.

## Functional consequences of pre- and postsynaptic STDP expression

The model reveals several functional implications of expressing synaptic plasticity pre- as well as postsynaptically. First, the locus of expression of plasticity will change the trial-to-trial variability of the synaptic response and overall reliability of neurotransmission. Specifically, by increasing the release probability, trial-to-trial reliability from synaptic transmission can be increased. Thus, joint pre- and postsynaptic plasticity can lead to a larger increase in the signal-to-noise ratio (SNR) than postsynaptic modification alone (Fig. 2a). The functional impact on SNR of this joint modification is consistent with improved sensory perception and its electrophysiological correlates observed in auditory cortex [97].

**Figure 2:**
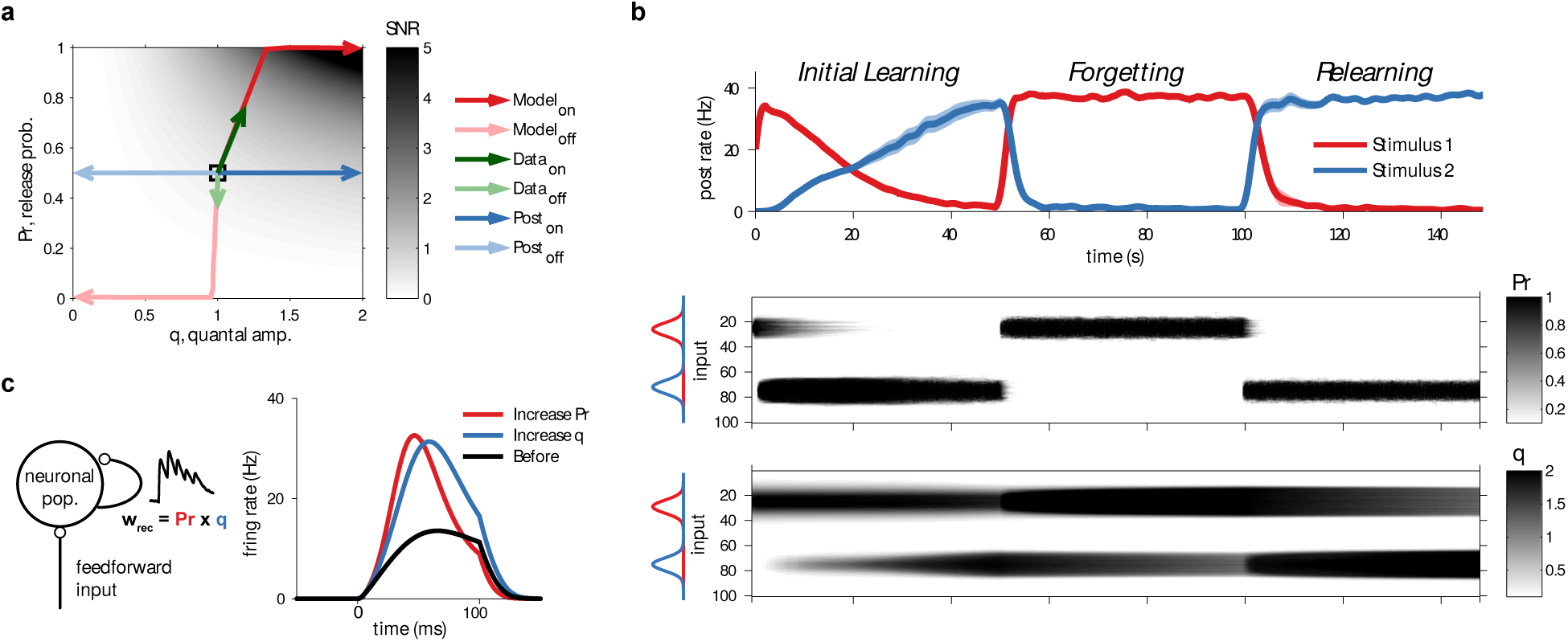
STDP with pre- and postsynaptic expression improves sensory perception, enables memory savings and shortens response latencies compared to postsynaptic expression alone. a) Changes in the signal-to-noise ratio (SNR) during receptive field learning in the STDP model. The SNR is represented by the gray-scale; the curves represent the various plasticity trajectories starting from the initial condition in the centre. Poisson train inputs that were stimulated at a high rate (“on”) obtain high signal-to-noise ratio (“SNR”) for postsynaptic-only potentiation (dark blue arrows), but combining pre- and postsynaptic potentiation yields considerably better SNR (dark red arrows). Weakly stimulated inputs (“off”) obtain lower SNR in either condition (light blue and light red arrows). These modelling results are in keeping with the observed modifications of *in-vivo* synaptic responses to a tone from on and off receptive field positions (dark and light green arrows) [97]. b) Rapid relearning and memory savings with asymmetrically combined pre- and postsynaptic expression of long-term plasticity. Top: Response of a neuron to two stimuli, red and blue. The neuron is initially trained on the blue stimulus, and becomes over time selective to it. This initial learning is slow because the changes in *q* (bottom panel) are slow. After learning, the memory is overwritten with the red stimulus. However, when switching back to the initial blue stimulus, the relearning is more rapid than at first exposure. Middle: Presynaptic LTP and LTD can rapidly completely reverse each other. Bottom: LTP has a postsynaptic component that does not reverse quickly, which means a postsynaptic trace is left behind after overwriting with novel information. This hidden trace enables rapid relearning of previously learnt, but overwritten, information. c) Left: Schematic of a firing-rate model with feedforward and feedback connections as described in [22]. In this network, recurrent synapses are short-term depressing. Changing release probability *Pr* affects the short-term dynamics, while changing the postsynaptic amplitude q only scales the postsynaptic response. Right: Comparison of changes in the response to a 100ms step stimulus in the recurrent network model when the recurrent synapses are subject to changes in either *Pr* or *q*. Increases in the release probability shorten the latency more than increases in the postsynaptic amplitude. Panels a and b were reproduced from [16].

Secondly, the pre- and postsynaptic components can differ in stability properties: some changes might be quick to induce, but hard to stabilise and vice versa. This in turn can provide neuronal networks with the necessary flexibility to quickly adapt to environmental changes. Using a simple receptive field development simulation, we propose that this might enable a form of memory savings. Memory savings is a concept introduced by Hermann Ebbinghaus and means that repeated learning of information is easier, even if the initially learned information appears to have been forgotten [98]. When memories were overwritten, the presynaptic component of the old memory was erased quickly but the postsynaptic component stayed largely intact. As a result, information that was initially learned but subsequently overwritten could rapidly be recovered upon relearning, provided that the postsynaptic component had not yet decayed completely (Fig. 2b). This mechanism could thus enable the brain to adapt quickly to different environments or to different tasks without fully forgetting previous learned information. The savings effect mirrors monocular deprivation experiments showing lasting postsynaptic structural effects on spine density that enable more rapid plasticity on repeated monocular deprivation [99, 100].

In the STDP data we saw no evidence for any decrease in the postsynaptic component *q*, perhaps because its decrease may be very slow. Under other protocols, LTD in q has been observed [68]. As it appears unbiological to have no decrease in *q*, we assumed that a slow homeostatic-like process can decrease *q* and so over very long times *q* decays and the hidden memory trace decays with it. Without this homeostatic process, the hidden trace in q would not decay and memory savings would occur for memories of any age. Our model also suggests that presynaptic boutons should be more dynamic during learning. Recently [101] imaged layer-5 pyramidal cell synapses and found that boutons tend to grow more often than spines after an auditory fear conditioning task.

Finally, while the effects reported in [16] considered feedforward networks, the changes in release probability under STDP also has consequences for recurrent networks. Excitation-dominated recurrent networks connected through strong short-term depressing synapses can have long response latencies, that are governed by the synaptic dynamics. We used the model presented in [22] to examine the effect of different expression loci in a recurrent network. Fig. 2c illustrates the response of a firing-rate model when the release probability *Pr* is increased, versus a case in which the quantal amplitude q is increased. The pre- and postsynaptic modifications were set such that the peak responses were identical. In both cases the response latency was shortened, but when release probability was allowed to increase due to LTP, response latency shortened about twice as much compared to the case where only postsynaptic plasticity was enabled.

## Possible other consequences of diversity in locus of plasticity

The “embarrassment of riches” in the possible expression sites of plasticity [47], is paralleled in many other biological systems. We mention the work of Eve Marder and co-workers on ion-channel expression [e.g. 102], and Turrigiano has emphasized the multiple ways to achieve homeostasis is puzzling (e.g. review Turrigiano in this issue). Considering Hebbian and homeostatic together (see Chen et al review in this issue), complicates this matter even further. It might have a number of consequences beyond the ones discussed above in the STDP model. First, the multiple expression site provide robustness to the system and multiple ways to maintain the capacity for plasticity, despite internal or external disruption, and compensate for genetic defects. Such redundancy can also be advantageous when an abundance of synapses is subject to somewhat diverse learning rules, as it increases the chance that one or some of the synapses correctly adapts to the task at hand. This diversity argument also occurs on the evolutionary level [103], namely, a population can be functionally similar but diverse in mechanism, allowing for better adaptation of the population as a whole to novel circumstances. Yet, the publication of yet another pathway often makes one want to exclaim “Who ordered that?”, as Rabi did when the sub-atomic muon particle was discovered.

Second, the multiple expression sites provide flexibility to local circuits, so that, via synapse-type-specific plasticity, different microcircuit components can be independently regulated [52]. For example, long-term depression (LTD) at layer 4 to layer 2/3 connections, but not at layer 2/3 to 2/3 synapses, is more readily induced during the critical period [104, 105], while thalamocortical LTP is already strongly diminished before the critical period has begun [106]. The locus of expression of long-term plasticity at these different synapse types also differs.

Similarly, different plasticity protocols are affected by distinct forms of neuromodulation. The neuromodulators can specifically control forms of STDP that express, for example, postsynaptically [107, 108, 109], providing a potential link between behaviourally relevant behaviours and expression loci.

Finally, LTD is not necessarily the opposite of LTP, this becomes even more pressing when considering the diversity of expression mechanisms. In virtually all computational models, LTP induction followed by LTD induction returns the synapse to its original state. Instead, in the above STDP model such a protocol might leave the synapse in a different state, even if the apparent synaptic weight is the same, as happens in the case of memory savings. A more direct experimental research of these issues, for instance using learning and subsequent unlearning, would be worthwhile. These considerations also indicates that both the pre- and postsynaptic component need mechanisms to prevent them from saturating and thereby losing the capacity for change. This might be possible by introducing soft-bounds for both the pre and post components, or introduce both pre and post synaptic normalization [110].

## Discussion

To model the impact of synaptic plasticity on circuit computations, it is important to know how synapses change during Hebbian and homoeostatic plasticity. Here, we have discussed several possible expression sites of synaptic plasticity. We have demonstrated three candidate effects in an STDP model where both pre- and postsynaptic components are modified: 1) a change in the release probability can improve the SNR in the circuit, 2) the difference in the time scales of modification can lead to the formation of hidden memory traces, and 3) as a result of changes in synaptic dynamics, the response latency in recurrent networks can be shortened with plasticity. The possible functional impact of combining pre- and postsynaptic plasticity is certainly not restricted to the three findings we illustrate here. We have rather just scratched the surface of what is likely an emerging field of study.

There is a large range of open issues. For instance, it has long been argued that the stability of memory in spite of continuous molecular turn-over is a quite remarkable problem for nature to solve [111, 112]. How synapses maintain stable information storage while staying plastic still remains unclear. The diversity of plasticity expression mechanisms could allow for a staged process by which initial changes are presynaptic, but later changes are consolidated structurally [32]. It is, however, not unlikely that multiple expression mechanisms are active in tandem. How these pre- and postsynaptic alterations are coordinated to ensure the long-term fidelity of information storage will require extensive further research. State-based models with a large range of transition rates between states have been explored to resolve this issue [113, 114, 115, 116], see also (Liu &Lisman, this issue). As these models are agnostic about expression, the current model could be seen as a biological implementation of such a multi-state model. It would for instance be of interest to know if the fast resetting of synaptic weights known to occur with exposure to enriched environments [117] is pre or post-synaptic. It would also be of interest to research if the storage capacity advantages observed in those more theoretical models will also occur in the current phenomenological model. There is also similarity to a recent study in which homeostasis acted as an independent multiplicative mechanism [118].

Another important issue is the weight dependence of long-term plasticity — LTP is hard to induce at synapses that are already strong [119, 120, 121, 93] — which has important implications for the synaptic weight distribution, memory stability [122] and information capacity [123]. It has been shown that presynaptic modifications strongly depend on the initial release probability [33], which is expected as release probability is bounded between 0 and 1. This demonstrates that the weight-dependence can stem from presynaptic considerations. However, postsynaptic mechanisms such as compartmentalisation of calcium signals may also explain this weight dependence, as it leads to large spines with long necks being “write protected” [124, 125, 126, 127]. This finding together with the fact that spine volume is proportional to the expression of AMPA receptors [128] implies that small spines should be more prone to LTP, which is consistent with experimental observations [69]. Such pre- and postsynaptic mechanisms are of course not mutually exclusive and both may contribute to the weight dependence of plasticity [120]. Including these effects would be an obvious next target for the STDP model. Experimentally, it would be of interest to apply protocols [see e.g. 87] that can accurately probe the short-term plasticity parameters before and after STDP induction.

Long-term synaptic plasticity and homeostatic plasticity have been fruitful modelling topics that have clarified the role of plasticity in biological neuronal networks as well as inspired applications using artificial neuronal networks. Yet, despite experimental evidence for presynaptic components in both Hebbian plasticity and synaptic homeostasis, in the overwhelming majority of computational models presynaptic contributions have been ignored (for an exception, see [129, 130]), or the models are agnostic about the expression and only adjust the synaptic weight. However, as we have seen, this is not a neutral assumption, and may affect the outcome of the plasticity on network function.

Interestingly, in recurrent networks short-term plasticity will have an effect on the pre/post activity patterns, and thereby change STDP induction [131, 132, 133]. Theoretically such mutually interacting systems are extremely challenging [134].

Our discussion has been restricted to the plasticity of excitatory synapses. Inhibitory neurons, in all their diversity [135, 136, 137], bring yet another level of complexity as differential short-term dynamics of excitatory and inhibitory synapses yields considerably richer dynamics [138, 139, 87, 62]. We suspect that only a small fraction of the richness and variety of the experimentally observed plasticity phenomena are understood and currently only a few computational models include them. A continued dialogue between theory and experiment should hopefully advance our understanding.

## Acknowledgements

We would like to thank our collaborators over the years for insightful discussions, in particular Rob Froemke, Richard Morris, Zahid Padamsey, Gina Turrigiano and Tim Vogels. We also like to acknowledge the workshop organisers Kevin Fox and Mike Stryker, and the Royal Society for facilitating an inspiring meeting.

## References

[1] Zhang, W. & Linden, D. J. The other side of the engram: experience-driven changes in neuronal intrinsic excitability. Nat. Rev. Neurosci. 4, 885–900 (2003).

[2] Watt, A. J. & Desai, N. S. Homeostatic plasticity and STDP: keeping a neuron’s cool in a fluctuating world. Front. Synaptic Neurosci. 2, 240 (2010).

[3] Keck, T. e. a. Integrating hebbian and homeostatic plasticity: the current state of the field and future research directions. Philosophical Transactions of the Royal Society B: Biological Sciences 369, 20130154 (2016).

[4] Poolos, N. P., Migliore, M. & Johnston, D. Pharmacological upregulation of h-channels reduces the excitability of pyramidal neuron dendrites. Nature neuroscience 5, 767–774 (2002).

[5] Hoffman, D. A., Magee, J. C., Colbert, C. M. & Johnston, D. *K*^+^ channel regulation of signal propagation in dendrites of hippocampal pyramidal neurons. Nature 387, 869–875 (1997).

[6] Desai, N. S., Rutherford, L. C. & Turrigiano, G. G. Plasticity in the intrinsic electrical properties of cortical pyramidal neurons. Nat. Neuro. 2, 515–520 (1999).

[7] Grubb, M. S. & Burrone, J. Activity-dependent relocation of the axon initial segment fine-tunes neuronal excitability. Nature 465, 1070–1074 (2010).

[8] Turrigiano, G. G. Too many cooks? intrinsic and synaptic homeostatic mechanisms in cortical circuit refinement. Annu. Rev. Neurosci. 34, 89–103 (2011).

[9] Turrigiano, G. The dialectic of hebb and homeostasis. Philosophical Transactions of the Royal Society B: Biological Sciences 369, 20130154 (2016).

[10] Markram, H. & Tsodyks, M. Redistribution of synaptic efficacy between neocortical pyramidal neurons. Nature 382, 807–810 (1996).

[11] Zakharenko, S., Zablow, L. & Siegelbaum, S. Visualization of changes in presynaptic function during long-term synaptic plasticity. Nature Neuroscience 4, 711–717 (2001).

[12] Zakharenko, S. S., Zablow, L. & Siegelbaum, S. A. Altered presynaptic vesicle release and cycling during mglur-dependent ltd. Neuron 35, 1099–1110 (2002).

[13] Zucker, R. S. & Regehr, W. G. Short-term synaptic plasticity. Annu Rev Physiol 64, 355–405 (2002).

[14] Gerdeman, G. L., Ronesi, J. & Lovinger, D. M. Postsynaptic endocannabinoid release is critical to long-term depression in the striatum. Nature neuroscience 5, 446–451 (2002).

[15] Sjöström, P. J., Turrigiano, G. G. & Nelson, S. B. Neocortical LTD via coincident activation of presynaptic NMDA and cannabinoid receptors. Neuron 39, 641–654 (2003).

[16] Costa, R. P., Froemke, R. C., Sjöström, P. J. & van Rossum, M. C. Unified pre-and postsynaptic long-term plasticity enables reliable and flexible learning. eLife 4, e09457 (2015).

[17] Abbott, L. F. & Regehr, W. G. Synaptic computation. Nature 431, 796–803 (2004).

[18] Hennig, M. H. Theoretical models of synaptic short term plasticity. Frontiers in Computational Neuroscience 7 (2013). URL http://www.frontiersin.org/computational_neuroscience/10.3389/fncom.2013.00045/abstract.

[19] Tsodyks, M. & Wu, S. Short-term synaptic plasticity. Scholarpedia (2013). revision #136920.

[20] Abbott, L. F., Varela, J. A., Sen, K. & Nelson, S. B. Synaptic depression and cortical gain control. Science 275, 220–224 (1997).

[21] Goldman, M. S., Maldonado, P. & Abbott, L. F. Redundancy reduction and sustained firing with stochastic depressing synapses. J Neurosci 22, 584–591 (2002).

[22] van Rossum, M. C. W., van der Meer, M. A. A., Xiao, D. & Oram, M. W. Adaptive integration in the visual cortex by depressing recurrent cortical circuits. Neural Comput 20, 1847–1872 (2008).

[23] Barak, O. & Tsodyks, M. Persistent activity in neural networks with dynamic synapses. PLoS Comput Biol 3, e35 (2007).

[24] York, L. C. & van Rossum, M. C. W. Recurrent networks with short term synaptic depression. J Comput Neurosci 27, 607–620 (2009).

[25] Mongillo, G., Barak, O. & Tsodyks, M. Synaptic theory of working memory. Science 319, 1543–1546 (2008).

[26] Markram, H., Gerstner, W. & Sjostrom, P. J. A history of spike-timing-dependent plasticity. Front Synaptic Neurosci 3, 4 (2011).

[27] Engert, F. & Bonhoeffer, T. Dendritic spine changes associated with hippocampal long-term synaptic plasticity. Nature 399, 66–70 (1999).

[28] Le Bé, J.-V. & Markram, H. Spontaneous and evoked synaptic rewiring in the neonatal neocortex. Proceedings of the National Academy of Sciences 103, 13214–13219 (2006).

[29] Kwon, H.-B. & Sabatini, B. L. Glutamate induces de novo growth of functional spines in developing cortex. Nature 474, 100–104 (2011).

[30] Nabavi, S. et al. Engineering a memory with LTD and LTP. Nature (2014).

[31] Kandel, E. R. The molecular biology of memory storage: a dialogue between genes and synapses. Science 294, 1030–1038 (2001).

[32] Loebel, A., Le Bé, J.-V., Richardson, M. J. E., Markram, H. & Herz, A. V. M. Matched pre- and post-synaptic changes underlie synaptic plasticity over long time scales. The Journal of neuroscience 33, 6257–6266 (2013).

[33] Larkman, A., Hannay, T., Stratford, K. & Jack, J. Presynaptic release probability influences the locus of long-term potentiation. Nature 360, 70–73 (1992).

[34] Bliss, T. V. & Collingridge, G. L. A synaptic model of memory: long-term potentiation in the hippocampus. Nature 361, 31–39 (1993).

[35] Younger, M. A., Müller, M., Tong, A., Pym, E. C. & Davis, G. W. A presynaptic ENaC channel drives homeostatic plasticity. Neuron 79, 1183–1196 (2013).

[36] Davis, G. W. & Müller, M. Homeostatic control of presynaptic neurotransmitter release. Annual review of physiology 77, 251–270 (2015).

[37] Kaeser, P. S. & Regehr, W. G. Molecular mechanisms for synchronous, asynchronous, and spontaneous neurotransmitter release. Annual review of physiology 76, 333 (2014).

[38] Müller, M., Liu, K. S. Y., Sigrist, S. J. & Davis, G. W. RIM controls homeostatic plasticity through modulation of the readily-releasable vesicle pool. The Journal of Neuroscience 32, 16574–16585 (2012).

[39] Wang, X., Pinter, M. J. & Rich, M. M. Reversible recruitment of a homeostatic reserve pool of synaptic vesicles underlies rapid homeostatic plasticity of quantal content. The Journal of Neuroscience 36, 828–836 (2016).

[40] Lee, H.-K., Barbarosie, M., Kameyama, K., Bear, M. F. & Huganir, R. L. Regulation of distinct AMPA receptor phosphorylation sites during bidirectional synaptic plasticity. Nature 405, 955–959 (2000).

[41] Kessels, H. W. & Malinow, R. Synaptic AMPA receptor plasticity and behavior. Neuron 61, 340–350 (2009).

[42] Isaac, J. T., Ashby, M. C. & McBain, C. J. The role of the GluR2 subunit in AMPA receptor function and synaptic plasticity. Neuron 54, 859–871 (2007).

[43] Raghavachari, S. & Lisman, J. E. Properties of quantal transmission at CA1 synapses. J Neurophysiol 92, 2456–2467 (2004).

[44] Lisman, J. E., Raghavachari, S. & Tsien, R. W. The sequence of events that underlie quantal transmission at central glutamatergic synapses. Nat Rev Neurosci 8, 597–609 (2007).

[45] Larkman, A. U. & Jack, J. J. B. Synaptic plasticity: hippocampal LTP. Current opinion in neurobiology 5, 324–334 (1995).

[46] Frey, U. & Morris, R. G. Synaptic tagging and long-term potentiation. Nature 385, 533–536 (1997).

[47] Malenka, R. C. & Bear, M. F. LTP and LTD: an embarrassment of riches. Neuron 44, 5–21 (2004).

[48] Sjöström, P. J., Turrigiano, G. G. & Nelson, S. Multiple forms of long-term plasticity at unitary neocortical layer 5 synapses. Neuropharmacology 52, 176–184 (2007).

[49] Kullmann, D. M., Moreau, A. W., Bakiri, Y. & Nicholson, E. Plasticity of inhibition. Neuron 75, 951–962 (2012).

[50] Padamsey, Z. & Emptage, N. Two sides to long-term potentiation: a view towards reconciliation. Philosophical Transactions of the Royal Society B: Biological Sciences 369, 20130154 (2014).

[51] Park, P. et al. Nmda receptor-dependent long-term potentiation comprises a family of temporally overlapping forms of synaptic plasticity that are induced by different patterns of stimulation. Phil. Trans. R. Soc. B 369, 20130131 (2014).

[52] Larsen, R. S. & Sjöström, P. J. Synapse-type-specific plasticity in local circuits. Current opinion in neurobiology 35, 127–135 (2015).

[53] Granger, A. J. & Nicoll, R. A. Expression mechanisms underlying long-term potentiation: a postsynaptic view, 10 years on. Phil. Trans. R. Soc. B 369, 20130136 (2014).

[54] Bacci, A. et al. Chronic blockade of glutamate receptors enhances presynaptic release and downregulates the interaction between synaptophysin-synaptobrevin-vesicle-associated membrane protein 2. J Neurosci 21, 6588–6596 (2001).

[55] Burrone, J., O’Byrne, M. & Murthy, V. N. Multiple forms of synaptic plasticity triggered by selective suppression of activity in individual neurons. Nature 420, 414–418 (2002). URL http://dx.doi.org/10.1038/nature01242.

[56] Thiagarajan, T. C., Lindskog, M. & Tsien, R. W. Adaptation to synaptic inactivity in hippocampal neurons. Neuron 47, 725–737 (2005). URL http://dx.doi.org/10.1016Zj.neuron.2005.06.037.

[57] Wierenga, C. J., Walsh, M. F. & Turrigiano, G. G. Temporal regulation of the expression locus of homeostatic plasticity. Journal of neurophysiology 96, 2127–2133 (2006).

[58] Han, E. B. & Stevens, C. F. Development regulates a switch between post- and presynaptic strengthening in response to activity deprivation. Proc Natl Acad Sci USA 106, 10817–10822. URL http://dx.doi.org/10.1073/pnas.0903603106.

[59] Kim, J. & Tsien, R. W. Synapse-specific adaptations to inactivity in hippocampal circuits achieve homeostatic gain control while dampening network reverberation. Neuron 58, 925–937 (2008).

[60] Ouanounou, G., Baux, G. & Bal, T. A novel synaptic plasticity rule explains homeostasis of neuromuscular transmission. eLife 5, e12190 (2016).

[61] Sanes, J. R. & Lichtman, J. W. Induction, assembly, maturation and maintenance of a postsynaptic apparatus. Nature Reviews Neuroscience 2, 791–805 (2001).

[62] Blackman, A. V., Abrahamsson, T., Costa, R. P., Lalanne, T. & Sjostrom, P. J. Targetcell-specific short-term plasticity in local circuits. Frontiers in Synaptic Neuroscience 5, 11 (2013).

[63] Markram, H., Wang, Y. & Tsodyks, M. Differential signaling via the same axon of neocortical pyramidal neurons. Proc Natl Acad Sci USA 95, 5323–5328 (1998).

[64] Buchanan, K. A. et al. Target-Specific Expression of Presynaptic NMDA Receptors in Neocortical Microcircuits. Neuron 75, 451–466 (2012).

[65] Koester, H. J. & Johnston, D. Target cell-dependent normalization of transmitter release at neocortical synapses. Science 308, 863–866 (2005).

[66] Banerjee, A., González-Rueda, A., Sampaio-Baptista, C., Paulsen, O. & Rodrífguez-Moreno, A. Distinct mechanisms of spike timing-dependent LTD at vertical and horizontal inputs onto L2/3 pyramidal neurons in mouse barrel cortex. Physiological reports (2014).

[67] Stanton, P. K., Winterer, J., Zhang, X.-l. & Müller, W. Imaging LTP of presynaptic release of FM1–43 from the rapidly recycling vesicle pool of Schaffer collateral-CA1 synapses in rat hippocampal slices. European Journal of Neuroscience 22, 2451–2461 (2005).

[68] Dodt, H.-U., Eder, M., Frick, A. & Zieglgänsberger, W. Precisely localized ltd in the neocortex revealed by infrared-guided laser stimulation. Science 286, 110–113 (1999).

[69] Matsuzaki, M., Honkura, N., Ellis-Davies, G. C. R. & Kasai, H. Structural basis of long-term potentiation in single dendritic spines. Nature 429, 761–766 (2004).

[70] Watt, A. J., Sjöström, P. J., Häusser, M., Nelson, S. B. & Turrigiano, G. G. A proportional but slower NMDA potentiation follows AMPA potentiation in LTP. Nature neuroscience 7, 518–524 (2004).

[71] Kullmann, D. M., Erdemli, G. & Asztély, F. LTP of AMPA and NMDA receptor-mediated signals: evidence for presynaptic expression and extrasynaptic glutamate spill-over. Neuron 17, 461–474 (1996).

[72] Hessler, N. A., Shirke, A. M. & Malinow, R. The probability of transmitter release at a mammalian central synapse (1993).

[73] Poncer, J. & Malinow, R. Postsynaptic conversion of silent synapses during LTP affects synaptic gain and transmission dynamics. Nature neuroscience 4, 989–996 (2001).

[74] Turrigiano, G. G., Leslie, K. R., Desai, N. S., Rutherford, L. C. & Nelson, S. B. Activity-dependent scaling of quantal amplitude in neocortical neurons. Nature 391, 892–896 (1998).

[75] Isaac, J., Oliet, S., Hjelmstad, G., Nicoll, R. & Malenka, R. Expression mechanisms of long-term potentiation in the hippocampus. Journal of Physiology-Paris 90, 299–303 (1996).

[76] Traynelis, S. F., Silver, R. A. & Cull-Candy, S. G. Estimated conductance of glutamate receptor channels activated during EPSCs at the cerebellar mossy fiber-granule cell synapse. Neuron 11, 279–289 (1993).

[77] Kilman, V., van Rossum, M. C. W. & Turrigiano, G. G. Activity deprivation reduces mIPSC amplitude by decreasing the number of postsynaptic GABAa receptors clustered at neocortical synapses. J. Neurosci. 22, 1328–1337 (2002).

[78] Feldwisch-Drentrup, H., Barrett, A. B., Smith, M. T. & van Rossum, M. C. Fluctuations in the open time of synaptic channels: An application to noise analysis based on charge. Journal of neuroscience methods 210, 15–21 (2012).

[79] Faber, D. S. & Korn, H. Applicability of the coefficient of variation method for analyzing synaptic plasticity. Biophysical journal 60, 1288–1294 (1991).

[80] Bolshakov, V. Y., Golan, H., Kandel, E. R. & Siegelbaum, S. A. Recruitment of new sites of synaptic transmission during the cAMP-dependent late phase of LTP at CA3-CA1 synapses in the hippocampus. Neuron 19, 635–651 (1997).

[81] Saez, I. & Friedlander, M. J. Plasticity between neuronal pairs in layer 4 of visual cortex varies with synapse state. The Journal of neuroscience 29, 15286–15298 (2009).

[82] Markram, H., L?bke, J., Frotscher, M., Roth, A. & Sakmann, B. Physiology and anatomy of synaptic connections between thick tufted pyramidal neurones in the developing rat neocortex. J Physiol 500 (Pt 2), 409–440 (1997).

[83] Loebel, A. et al. Multiquantal release underlies the distribution of synaptic efficacies in the neocortex. Frontiers in computational neuroscience 3, 27 (2009).

[84] Rozov, A. & Burnashev, N. Polyamine-dependent facilitation of postsynaptic ampa receptors counteracts paired-pulse depression. Nature 401, 594–598 (1999).

[85] Rozov, A., Jerecic, J., Sakmann, B. & Burnashev, N. AMPA receptor channels with long-lasting desensitization in bipolar interneurons contribute to synaptic depression in a novel feedback circuit in layer 2/3 of rat neocortex. The Journal of Neuroscience 21, 8062–8071 (2001).

[86] Heine, M. et al. Surface mobility of postsynaptic AMPARs tunes synaptic transmission. Science 320, 201–205 (2008).

[87] Costa, R. P., Sjostrom, P. J. & van Rossum, M. C. W. Probabilistic inference of short-term synaptic plasticity in neocortical microcircuits. Frontiers in Computational Neuroscience 7, 75 (2013).

[88] Bhumbra, G. S. & Beato, M. Reliable evaluation of the quantal determinants of synaptic efficacy using Bayesian analysis. Journal of Neurophysiology 109, 603–620 (2013).

[89] Barri, A., Wang, Y., Hansel, D. & Mongillo, G. Quantifying repetitive transmission at chemical synapses: A generative-model approach. eNeuro 3, ENEURO-0113 (2016).

[90] Bender, V. A. & Feldman, D. Two Coincidence Detectors for Spike Timing-Dependent Plasticity in Somatosensory Cortex. The Journal of neuroscience 26, 4166–4177 (2006).

[91] Yang, Y. & Calakos, N. Presynaptic long-term plasticity. Frontiers in Synaptic Neuroscienc 5 (2013).

[92] Andrade-Talavera, Y., Duque-Feria, P., Paulsen, O. & Rodrífguez-Moreno, A. Presynaptic spike timing-dependent long-term depression in the mouse hippocampus. Cerebral Cortex 26, 3637–3654 (2016).

[93] Sjöström, P. J., Turrigiano, G. G. & Nelson, S. B. Rate, Timing, and Cooperativity Jointly Determine Cortical Synaptic Plasticity. Neuron 32, 1149–1164 (2001).

[94] Pfister, J.-P. & Gerstner, W. Triplets of spikes in a model of spike timing-dependent plasticity. J Neurosci 26, 9673–9682 (2006).

[95] Clopath, C., Büsing, L., Vasilaki, E. & Gerstner, W. Connectivity reflects coding: a model of voltage-based stdp with homeostasis. Nat Neurosci 13, 344–352 (2010).

[96] Graupner, M. & Brunel, N. Calcium-based plasticity model explains sensitivity of synaptic changes to spike pattern, rate, and dendritic location. Proc Natl Acad Sci USA 109, 3991–3996 (2012).

[97] Froemke, R. C. et al. Long-term modification of cortical synapses improves sensory perception. Nature Neuroscience 16, 79–88 (2013).

[98] Ebbinghaus, H. Memory: A contribution to experimental psychology. Teachers College, Columbia University (1913).

[99] Hofer, S., Mrsic-Flogel, T., Bonhoeffer, T. & Hübener, M. Experience leaves a lasting structural trace in cortical circuits. Nature 457, 313–317 (2008).

[100] Hübener, M. & Bonhoeffer, T. Searching for Engrams. Neuron 67, 363–371 (2010).

[101] Yang, Y. et al. Selective synaptic remodeling of amygdalocortical connections associated with fear memory. Nature Publishing Group 19, 1348–1355 (2016).

[102] Marder, E. & Goaillard, J.-M. Variability, compensation and homeostasis in neuron and network function. Nature Reviews Neuroscience 7, 563–574 (2006).

[103] Daniels, B. C., Chen, Y.-J., Sethna, J. P., Gutenkunst, R. N. & Myers, C. R. Sloppiness, robustness, and evolvability in systems biology. Current Opinion in Biotechnology 19, 389–395 (2008).

[104] Corlew, R., Wang, Y., Ghermazien, H., Erisir, A. & Philpot, B. D. Developmental switch in the contribution of presynaptic and postsynaptic NMDA receptors to long-term depression. The Journal of Neuroscience 27, 9835–9845 (2007).

[105] Banerjee, A. et al. Double dissociation of spike timing-dependent potentiation and depression by subunit-preferring NMDA receptor antagonists in mouse barrel cortex. Cerebral Cortex 19, 2959–2969 (2009).

[106] Crair, M. C. & Malenka, R. C. A critical period for long-term potentiation at thalamocortical synapses. Nature 375, 325–328 (1995).

[107] Seol, G. H. et al. Neuromodulators control the polarity of spike-timing-dependent synaptic plasticity. Neuron 55, 919–929 (2007).

[108] Pawlak, V., Wickens, J. R., Kirkwood, A. & Kerr, J. N. D. Timing is not everything: neuromodulation opens the STDP gate. Frontiers in Synaptic Neuroscience (2010).

[109] Frémaux, N. & Gerstner, W. Neuromodulated Spike-Timing-Dependent Plasticity, and Theory of Three-Factor Learning Rules. Frontiers in Neural Circuits 9, 1178 (2016).

[110] Zenke, F. & Gerstner, W. Hebbian plasticity requires negative feedback on multiple timescales. Philosophical Transactions of the Royal Society B: Biological Sciences 369, 20130154 (2016).

[111] Crick, F. Memory and molecular turnover. Nature 312, 101 (1984).

[112] Lisman, J. A mechanism for the hebb and the anti-hebb processes underlying learning and memory. Proceedings of the National Academy of Sciences 86, 9574–9578 (1989).

[113] Fusi, S., Drew, P. J. & Abbott, L. Cascade Models of Synaptically Stored Memoriest. Neuron 45, 599–611 (2005). URL http://dx.doi.org/ron.2005.02.001.

[114] Leibold, C. & Kempter, R. Sparseness constrains the prolongation of memory lifetime via synaptic metaplasticity. Cerebral Cortex 18, 67–77 (2008).

[115] Barrett, A. B. & van Rossum, M. C. W. Optimal learning rules for discrete synapses. PLoS Comput Biol 4, e1000230 (2008).

[116] Lahiri, S. & Ganguli, S. A memory frontier for complex synapses. In Advances in Neural Information Processing Systems, 1034–1042 (2013).

[117] Abraham, W. C., Logan, B., Greenwood, J. M. & Dragunow, M. Induction and Experience-Dependent Consolidation of Stable Long-Term Potentiation Lasting Months in the Hippocampus. J. Neurosci. 22, 9626–9634 (2002).

[118] Toyoizumi, T., Kaneko, M., Stryker, M. P. & Miller, K. D. Modeling the dynamic interaction of hebbian and homeostatic plasticity. Neuron 84, 497–510 (2014). URL http://dx.doi.org/10.1016/j.neuron.2014.09.036.

[119] Bi, G.-q. & Poo, M.-m. Synaptic modifications in cultured hippocampal neurons: dependence on spike timing, synaptic strength, and postsynaptic cell type. J. Neurosci. 18, 10464–10472 (1998).

[120] Liao, D., Jones, A. & Malinow, R. Direct measurement of quantal changes underlying long-term potentiation in ca1 hippocampus. Neuron 9, 1089–1097 (1992).

[121] Debanne, D., Gähwiler, B. H. & Thompson, S. M. Heterogeneity of synaptic plasticity at unitary CA1-CA3 and CA3-CA3 connections in rat hippocampal slice cultures. J. Neurosci. 19, 10664–10671 (1999).

[122] Billings, G. & van Rossum, M. C. W. Memory retention and spike-timing-dependent plasticity. J Neurophysiol 101, 2775–2788 (2009).

[123] van Rossum, M., Shippi, M. & Barrett, A. Soft-bound synaptic plasticity outperforms hardbound plasticity (2012). Submitted PlosCB.

[124] Noguchi, J., Matsuzaki, M., Ellis-Davies, G. C. R. & Kasai, H. Spine-neck geometry determines NMDA receptor-dependent Ca2+ signaling in dendrites. Neuron 46, 609–622 (2005).

[125] Yasumatsu, N., Matsuzaki, M., Miyazaki, T., Noguchi, J. & Kasai, H. Principles of long-term dynamics of dendritic spines. J Neurosci 28, 13592–13608 (2008). URL http://dx.doi.org/10.1523/JNEUR0SCI.0603–08.2008.

[126] Kalantzis, G. & Shouval, H. Z. Structural plasticity can produce metaplasticity. PLoS One 4, e8062 (2009).

[127] O’Donnell, C., Nolan, M. F. & van Rossum, M. C. W. Dendritic spine dynamics regulate the long-term stability of synaptic plasticity. J Neurosci 31, 16142–16156 (2011).

[128] Matsuzaki, M. et al. Dendritic spine geometry is critical for AMPA receptor expression in hippocampal CA1 pyramidal neurons. Nature neuroscience 4, 1086–1092 (2001).

[129] Senn, W., Markram, H. & Tsodyks, M. An algorithm for modifying neurotransmitter release probability based on pre-and postsynaptic spike timing. Neural Computation 13, 35–67 (2001).

[130] Lisman, J. & Raghavachari, S. A unified model of the presynaptic and postsynaptic changes during LTP at CA1 synapses. Sci STKE 2006, re11 (2006).

[131] Wang, Y. et al. Heterogeneity in the pyramidal network of the medial prefrontal cortex. Nat Neurosci 9, 534–542 (2006).

[132] Vasilaki, E. & Giugliano, M. Emergence of Connectivity Motifs in Networks of Model Neurons with Short- and Long-Term Plastic Synapses. PLoS ONE 9, e84626 (2014).

[133] Froemke, R. & Dan, Y. Spike-timing-dependent synaptic modification induced by natural spike trains. Nature 416, 433–8 (2002).

[134] Gilson, M., Burkitt, A., Grayden, D., Thomas, D. & van Hemmen, J. Emergence of network structure due to spike-timing-dependent plasticity in recurrent neuronal networks. I. input selectivity-strengthening correlated input pathways. Biological cybernetics 101, 81–102 (2009).

[135] Markram, H. et al. Interneurons of the neocortical inhibitory system. Nature Reviews Neuroscience 5, 793–807 (2004).

[136] DeFelipe, J. et al. New insights into the classification and nomenclature of cortical gabaergic interneurons. Nature Reviews Neuroscience 14, 202–216 (2013).

[137] Kepecs, A. & Fishell, G. Interneuron cell types are fit to function. Nature 505, 318–326 (2014).

[138] Galarreta, M. & Hestrin, S. Frequency-dependent synaptic depression and the balance of excitation and inhibition in the neocortex. Nature neuroscience 1, 587–594 (1998).

[139] Varela, J. A., Song, S., Turrigiano, G. G. & Nelson, S. B. Differential depression at excitatory and inhibitory synapses in visual cortex. The journal of neuroscience 19, 4293–4304 (1999).

